# VDR-calcium axis regulates the diet-driven metabolic shift during weaning

**DOI:** 10.1101/2025.01.16.633313

**Authors:** Neha Jawla, Shubhi Khare, Nidhi Yadav, Ranjan Kumar Nanda, Gopalakrishnan Aneeshkumar Arimbasseri

**Author notes:** These authors contributed equally (Neha Jawla, Shubhi Khare).

## Abstract

Weaning in mammals is associated with a shift in the metabolism, driven by the differences in the macronutrient composition of milk and post-weaning diet. Milk has a higher fat content compared with the carbohydrate-enriched solid food. Malnutrition during this stage could affect this transition with long-term adverse effects. The role of micronutrients during this transition is not well understood. Mice lacking a functional vitamin D receptor (VDR) progressively develop severe skeletal muscle and adipose atrophy after weaning, suggesting a role for vitamin D signaling in the metabolic transition during weaning. Here, we demonstrate that after weaning, VDR knock-out mice exhibit systemic energy deprivation and higher lipolysis in inguinal white adipose tissue, probably due to increased norepinephrine signaling via protein kinase A (PKA) and extracellular signalling-regulated kinase (ERK) pathways. Energy deprivation in *vdr*^-/-^ mice is associated with defective liver glycogenolysis, characterized by increased expression of protein phosphatase-1α and decreased phosphorylation of glycogen phosphorylase. However, restoration of serum calcium levels by a rescue diet is sufficient to restore energy metabolism in *vdr*^-/-^ mice. Interestingly, maintaining a high-fat-containing milk-based diet post-weaning could prevent the onset of energy deprivation, liver glycogen storage defect, and adipose atrophy in these mice without restoring serum calcium levels. Our data show that the vitamin D-calcium axis is essential for the adaptation of mice to the dietary shift from high-fat-containing milk to post-weaning carbohydrate-enriched diets. It also reveals a novel macronutrient-micronutrient interaction that shapes the metabolic flexibility of the individual based on the dietary composition of nutrients.

## Introduction

Weaning involves a dietary switch in mammals, progressively decreasing the reliance on the consumption of a fat-rich milk diet in favor of a carbohydrate-rich diet (1, 2). Metabolic adaptation to this shift in macronutrient consumption is characterized by reduced hepatic gluconeogenesis, increased liver glycogen content, and changes in lipid metabolism (2). Such metabolic changes are supported by various nutritional, hormonal, and neuronal factors (3, 4). Dietary changes during weaning are shown to drive β-cell proliferation and maturation, which is important for the optimal endocrine function of the pancreas (5). A switch from the nutrient sensor target of rapamycin (mTORC1) to the energy sensor 5’-adenosine monophosphate-activated protein kinase (AMPK) was found critical for functional maturation of β-cells (6). Furthermore, changes in the macronutrient composition during the weaning process drive alterations in the gut microbiome, which is essential for the development of immune tolerance (7). The major calcium absorption pathway also changes during weaning, from the paracellular pathway during the suckling stage to the vitamin D dependent transcellular pathway post-weaning (8). However, the factors that regulate these post-weaning metabolic adaptations are not fully understood.

Vitamin D is a steroid hormone, and its biologically active form (1, 25(OH)_2_D) mediates its genomic effects by binding to an intracellular receptor called vitamin D receptor (9). VDR regulates several genes involved in energy balance, glucose homeostasis, and hormone regulation (10). In mice, the absence of VDR signaling has been shown to lead to growth retardation, hypocalcemia, hypophosphatemia, hyperparathyroidism, and rickets after weaning. These mice, however, despite a lack of VDR, show no signs of morphological or metabolic abnormalities and grow normally throughout early development until weaning (11, 12). *vdr*^-/-^ mice recapitulate abnormalities observed in humans with vitamin D dependent rickets type II (VDDR II), caused by inactivating mutations in VDR (13). Earlier, we showed that maintaining *vdr*^-/-^ mice on high-fat-containing milk mimetic diets prevented skeletal muscle abnormalities and rescued pancreatic insulin response (14). These observations raise the question of whether the dietary transition from high-fat milk to high-carbohydrate chow is dependent on VDR signaling.

Vitamin D signaling is essential for mineral homeostasis by controlling intestinal calcium absorption, renal calcium re-absorption, and calcium resorption from the bones (8, 15). Restoration of intestinal VDR in whole-body VDR-knockout mice resulted in normocalcemia, and was sufficient to recover from the growth and skeletal phenotypes (16). A calcium and phosphate enriched; lactose-supplemented diet was also shown to prevent the development of growth and bone abnormalities in *vdr*^-/-^ mice (17). These data raise the question of whether the vitamin D-calcium axis plays any role in the metabolic shifts happening during the weaning process.

Previous observations from our lab have shown that vitamin D receptor knockout mice develop severe skeletal muscle atrophy associated with increased protein degradation and severe energy deprivation because of a glycogen storage defect (18). However, this does not explain the systemic defects in glucose homeostasis in *vdr*^-/-^ mice. In the present study, we find that systemic loss of VDR signaling leads to glycogen utilization defect in the liver, associated with lower activity of the liver isoform of glycogen phosphorylase (PYGL) and increased levels of protein phosphatase 1. As observed in the genetic disorder caused by glycogen phosphorylase deficiency, we observed hypoglycemia in these mice. This systemic energy deprivation caused increased norepinephrine levels in the adipose tissue, leading to activation of protein kinase A and ERK pathways. These, in turn, activate hormone-sensitive lipase, culminating in adipose atrophy and the establishment of cachexia. Normalizing serum calcium by providing a calcium-rich rescue diet negates the effects of VDR ablation on liver glycogenolysis, adipose wasting, and energy balance. More importantly, we show that the absence of VDR signaling can be compensated by maintaining the mice on a high-fat diet without the need to normalize serum calcium. Together, this supports the important role of the vitamin D-calcium axis in orchestrating the shift from fat to carbohydrate metabolism.

## Results

### *vdr*-/-mice exhibit severe energy deprivation and adipose atrophy post-weaning

We earlier showed that vitamin D receptor knockout (*vdr*^-/-^) mice exhibit severe muscle wasting associated with muscle glycogen accumulation after weaning (18), and milk-based, fat-enriched diets could rescue muscle mass in these mice (14). We hypothesized that their inability to utilize carbohydrates as a primary energy source after weaning leads to systemic energy deprivation, causing severe growth defects and wasting phenotype.

In support of systemic energy deficiency, blood glucose levels in *vdr*^-/-^ mice were low compared to wild type (WT) mice under both fed and fast conditions (Figure 1A&B). Furthermore, the glucagon:insulin ratio was also higher in the *vdr*^-/-^ mice (Figure 1C). Blood leptin concentrations, a sensor of the energy reservoir in the body (19, 20) were also dramatically reduced in *vdr*^-/-^ mice under *ad libitum* feeding conditions (Figure 1D). There was no change in serum levels of ghrelin, an appetizing hormone, between WT and *vdr*^-/-^ mice (Figure S1A).

**Fig. 1.**
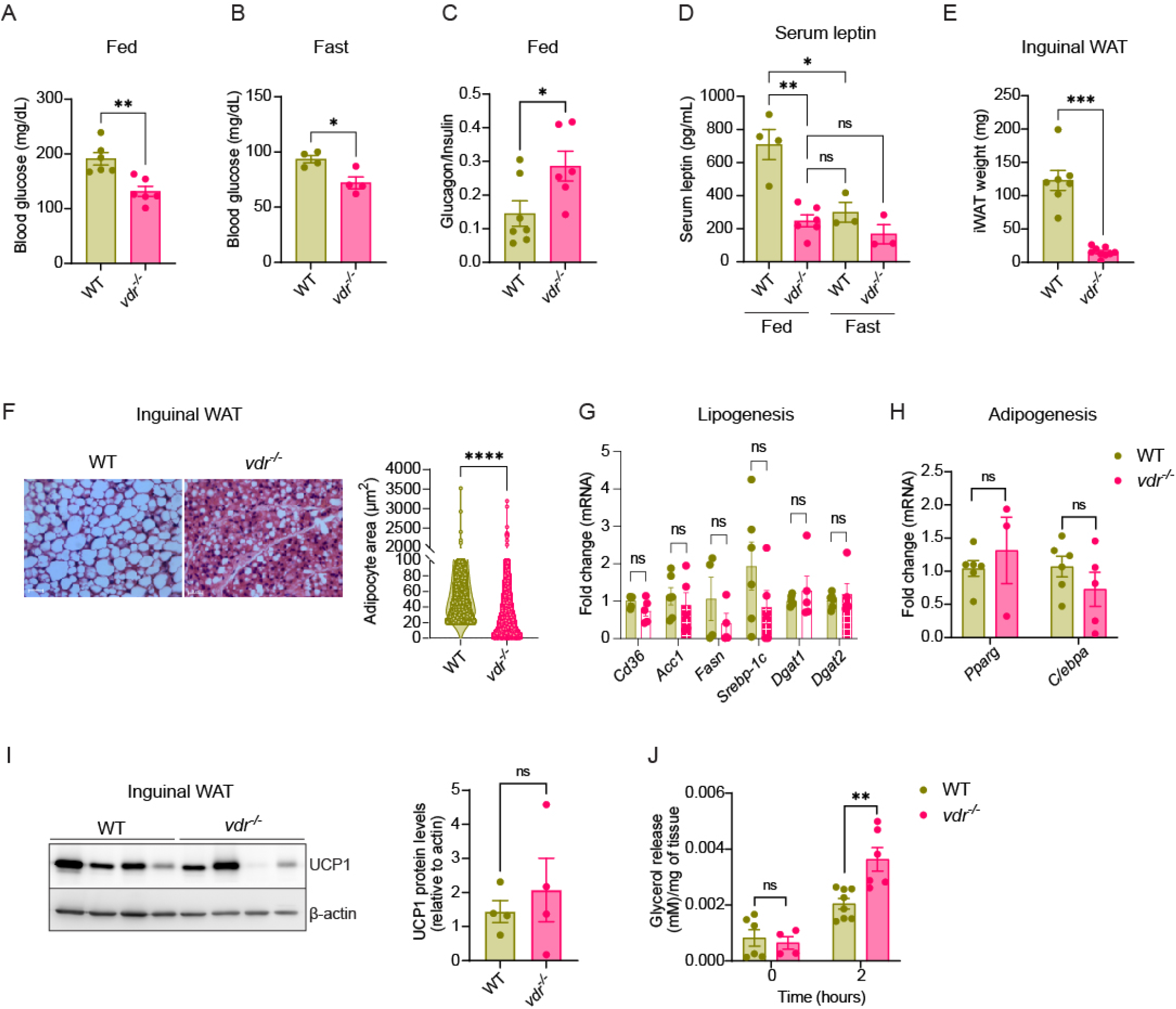
*vdr* ^-/-^ mice exhibit severe energy deprivation and adipose atrophy post-weaning. (A&B) Blood glucose levels in *ad libitum* fed state (A) and after 12 hours of fasting (B) in 7-week WT and *vdr* ^-/-^ mice. (C) Serum glucagon/insulin ratio in 7-week WT and *vdr* ^-/-^ mice under *ad libitum* fed state measured using ELISA. (D) Serum levels of leptin in 7-week WT and *vdr* ^-/-^ mice under *ad libitum* fed and fast state (12 hours) measured using ELISA. (E) Inguinal WAT weight (in mg) of 7-week WT and *vdr* ^-/-^ mice. (F) Representative images of H&E stained iWAT sections and quantitative analysis of adipocyte area in 7-week WT and *vdr* ^-/-^ mice (right panel). (G&H) mRNA expression levels of genes involved in lipogenesis (G) and adipogenesis (H) in iWAT of 7-week WT and *vdr* ^-/-^ mice as quantified using qRT-PCR. (I) Representative western blot and accompanied densitometric quantification for UCP-1 in iWAT of 7-week WT and *vdr* ^-/-^ mice. (J) Glycerol release from iWAT explants of 7-week WT and *vdr* ^-/-^ mice. All graphs show mean ± SEM. Statistical significance was determined by unpaired t-test (*p < 0.05, **p < 0.01, ***p< 0.001 and ****p < 0.0001). The number of samples is denoted by the dots in the graphs.

Adipose tissue shows remarkable plasticity undergoing hyperplasia, hypertrophy, and shrinkage based on the metabolic requirement of the body (21). We observed drastically reduced inguinal white adipose tissue weight (iWAT) in *vdr*^-/-^ mice compared to WT mice (Figure 1E). In line with previous reports (22), histopathological analysis of cross-sections of inguinal white adipose tissue shows reduced adipocyte size in these mice (Figure 1F). However, no discernible difference in adipose weight and adipocyte area was observed at 3 weeks of age when *vdr*^-/-^ mice were indistinguishable from WT mice (Figure S1B&C). Accordingly, leptin and glucose levels were comparable at this stage between WT and *vdr*^-/-^ mice (Figure S1D&E). These results clearly indicate that the energy deprivation in *vdr*^-/-^ mice originates post-weaning.

Next, we asked if the adipose wasting in *vdr*^-/-^ is indicative of defective adipogenesis and lipogenesis. To address this, we performed qRT-PCR analysis of the genes involved in these pathways in iWAT of 7-week-old WT and *vdr*^-/-^ mice. Intriguingly, expression levels of genes involved in lipogenesis (*Cd36, Acc1, Fasn, Srebp1, Dgat1&2*) and adipogenesis (*Pparg & C/ebpa*) were unchanged in the iWAT of these mice (Figure 1G&H) suggesting that adipose wasting in *vdr*^-/-^ mice is not due to defects in fatty acid synthesis or adipose development. Furthermore, neither the protein levels nor the phosphorylation of ACC1, a key enzyme in the fatty acid biosynthetic pathway, showed any difference between WT and *vdr*^-/-^ (Figure S1F). We also checked if *vdr*^-/-^ exhibits increased energy expenditure via enhanced thermogenesis (22, 23). However, we did not observe any consistent difference in uncoupling protein 1 (UCP1) level, a mitochondrial protein that uncouples oxidative phosphorylation from ATP synthesis facilitating energy dissipation, in iWAT of *vdr*^-/-^ mice (Figure 1I). Transcript levels of *Ucp1* and *Ucp2* also did not differ between WT and *vdr*^-/-^ adipose (Figure S1G&H). These results suggest that the adipose wasting in the *vdr*^-/-^ mice was not due to an increase in thermogenesis or defect in adipogenesis or lipogenesis.

Negative energy balance is considered one of the primary physiological cues to initiate lipolysis to increase fat mobilization from adipose depot to maintain the energy requirements of the body (20). To check if lipolysis in iWAT is higher in the *vdr*^-/-^ mice, we analyzed glycerol release, an indicator of lipolysis, by adipose explants *ex vivo* (24). Higher glycerol release from adipose explants of 7-week-old *vdr*^-/-^ was detected within 2 hours of incubation (Figure 1J). These data demonstrate that *vdr*^-/-^ mice display adipose atrophy characterized by enhanced basal lipolysis.

### *vdr*-/-mice display increased norepinephrine levels with increased PKA and ERK activity in iWAT

The lipolysis program of adipose tissue is activated by several signals, including glucagon and catecholamines. These pathways work through the activation of protein kinase A (PKA), a downstream modulator of the β-adrenergic signaling pathway, and subsequent phosphorylation of hormone-sensitive lipase (25, 26). We observed that the PKA activity was upregulated in the *vdr*^-/-^ mice (Figure 2A). Moreover, the total and phosphorylated form (Ser 563) of HSL was also high in the *vdr*^-/-^ adipose (Figure 2B). These results clearly indicate that increased PKA and HSL activities drive severe adipose atrophy in *vdr*^-/-^ mice. Insulin signaling is known to restrain lipolysis via Akt-mediated phosphodiesterase 3b (PDE3B) activation, which in turn leads to a reduction in PKA activity (27, 28). We reasoned that lower levels of insulin in *vdr*^-/-^ might contribute to enhanced lipolysis (18). However, phosphorylation of Akt (Ser 473), an effector of the insulin signaling pathway, was unaffected in adipose of *vdr*^-/-^ and WT mice (Figure S2A). Moreover, the activity of the mTORC1 pathway, another suppressor of lipolysis, as measured by the levels of p-S6, was also unchanged in *vdr*^-/-^ adipose (Figure S2B).

**Fig. 2.**
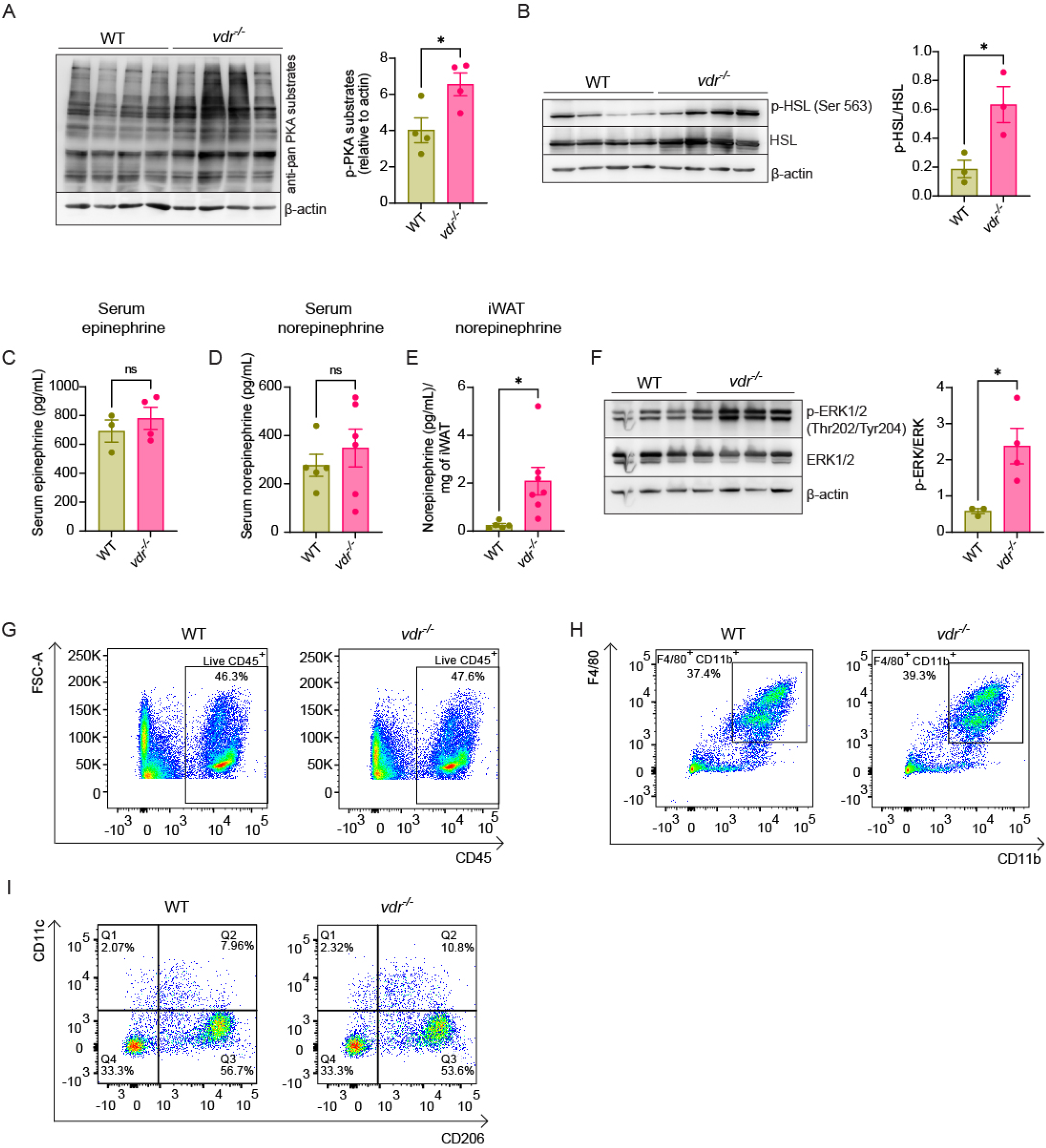
*vdr* ^-/-^ mice display increased norepinephrine levels with increased PKA and ERK activity in iWAT. (A) Representative western blot and densitometric quantification of phospho-PKA substrates (RRXS*/T*), normalized against β-actin in iWAT of 7-week WT and *vdr* ^-/-^ mice. (B) Representative images of western blot showing phospho-HSL (Ser 563), HSL, β-actin, and the ratio of phospho-HSL to total HSL (right panel) in iWAT of 7-week WT and *vdr* ^-/-^ mice. (C&D) Serum levels of epinephrine (C) and norepinephrine (D) in 7-week WT and *vdr* ^-/-^ mice, measured using ELISA. (E) Norepinephrine levels in iWAT of 7-week WT and *vdr* ^-/-^ mice, measured using ELISA. (F) Western blot representing phospho-ERK1/2 (Thr 202/ Tyr 204), ERK, and β-actin accompanied with the ratio of phospho-ERK 1/2 to total ERK1/2 in iWAT of 7-week WT and *vdr* ^-/-^ mice. (G-I) Immune profiling in iWAT of 7-week WT and *vdr* ^-/-^ mice. Representative FACS plots of live CD45^+^ cells (G), F4/80 versus CD11b (H) and CD11c versus CD206 (I). The number within the plots indicates the percentage of cells. Quantification of the same is given in Figure S2 (D-F). All graphs show mean ± SEM. Statistical significance was determined by unpaired t-test (*p < 0.05, **p < 0.01, ***p< 0.001 and ****p < 0.0001). The number of samples is denoted by the dots in the graphs.

Since catecholamine-mediated activation of β-adrenergic receptors constitutes one of the major regulators of the prolipolytic PKA pathway, we measured the level of serum epinephrine and norepinephrine. However, the serum levels of both hormones were unchanged in *vdr*^-/-^ mice (Figure 2C&D). The transcript levels of β-adrenergic receptors in iWAT of *vdr*^-/-^ mice were not different from those of WT (Figure S2C). Studies have shown that white adipose tissue is abundantly innervated by sympathetic nerves that release norepinephrine in the local environment (29). We, therefore, measured the norepinephrine concentrations within iWAT. Adipose tissue from *vdr*^-/-^ mice showed a remarkable increase in norepinephrine levels compared to WT (Figure 2E), suggesting increased catecholamine levels in the adipose tissue.

Apart from the well-established beta-adrenergic signaling, extracellular signal-regulated kinase proteins (ERK1/2) can drive lipolysis independently as well as in combination with β-adrenergic signalling (30, 31). Recent studies have shown evidence of the involvement of activated ERK in increased basal lipolysis in adipose tissue of obese mice (32). When compared with WT mice, total ERK protein levels were unchanged in *vdr*^-/-^ mice; on the contrary, the levels of ERK1/2 phosphorylated at Thr202/Tyr204 were significantly increased in *vdr*^-/-^ adipose, confirming the activation of the ERK signaling pathway (Figure 2F). ERK activity is shown to be induced via inflammatory cytokines (33), and vitamin D is known to have anti-inflammatory activity (34, 35). Increased lipolysis has been associated with increased macrophage infiltration and inflammation (32). We found no differences in the relative proportion of CD45 positive cells between the control and *vdr*^-/-^ iWAT (Figure 2G & S2D). Additionally, the relative levels of CD11b^+^F4/80^+^ macrophages were also unchanged in *vdr*^-/-^ adipose tissue (Figure 2H & S2E). The polarisation status of macrophages regulates the balance between pro and anti-inflammatory milieu within adipose (36), with several studies highlighting a shift towards CD11c^+^ M1 macrophage in obese conditions, whereas M2 (CD206^+^) macrophage predominates in lean individuals (37). We observed a higher proportion of M2 (CD11c^-^CD206^+^) macrophages in both WT and *vdr*^-/-^ mice (Figure 2I & S2F). Furthermore, there was no difference in the mRNA levels of cytokines *Tnfa, IL-6, Ifng*, and *Il10* in adipose of *vdr*^-/-^ mice compared to WT (Figure S2G). Inflammation-mediated signaling pathways engage the induction of NF-𝓍B, which is associated with I𝓍Bα degradation, a potent NF-𝓍B inhibitor (38, 39). Total protein levels of I𝓍Bα were similar between WT and *vdr*^-/-^ adipose (Figure S2H), a further indication that adipose atrophy in *vdr*^-/-^ mice is not induced by inflammatory signals. Collectively, these findings suggest that adipose β3-adrenergic receptor stimulation-induced lipolysis is associated with increased norepinephrine levels in adipose, leading to adipose wasting in *vdr*^-/-^ mice.

### *vdr*-/-mice show increased adiponectin and enhanced adipose mitochondrial activity

A previous study from our lab showed that ablation of VDR signalling leads to severe energy deprivation in skeletal muscles (18). To understand if other metabolic tissues like the liver and adipose also exhibit energy imbalance in the absence of vitamin D signaling, we measured ATP levels in the liver and adipose. Similar to skeletal muscles, liver from *vdr*^-/-^ mice showed energy insufficiency reflected in significantly lower ATP levels (Figure 3A). On the contrary, ATP levels were maintained in *vdr*^-/-^ adipose (Figure 3B). This is in line with increased PKA activity in adipose tissue that has been shown to enhance mitochondrial metabolism (40). To gain further insight into the underlying mechanisms of how adipose tissue maintains its energy levels despite energy deficiency in other metabolic tissues, we analysed mitochondrial mass and function. The expression of TOMM20, a central component of the mitochondrial protein import machinery located on the outer membrane was augmented in *vdr*^-/-^ adipose tissue compared to WT indicating increased mitochondrial mass (Figure 3C). To further corroborate our findings, we examined adipose peroxisome proliferator-activated receptor gamma coactivator 1-α (PGC1α) expression, a well-characterized indicator of mitochondrial biogenesis. mRNA levels of *Ppargc1a* (PGC1α) were significantly increased in *vdr*^-/-^ adipose (Figure 3D). PGC1α expression is directly regulated by cyclic AMP response element binding (CREB) protein, one of the targets of PKA (41). *vdr*^-/-^ adipose showed higher CREB activity as indicated by increased phosphorylation of CREB at Ser 133 (Figure S3A). However, western blot analysis of the relative levels of mitochondrial and nuclear-encoded proteins involved in OXPHOS showed subunits of complex V (ATP5A), complex II (SDHB), complex III (UQCRC2), complex IV (COX II), and complex I (NDUFB8) were similar in the *vdr*^-/-^ & WT adipose (Figure 3E). We then evaluated metabolic function directly using high-resolution tissue respirometry and found that *vdr*^-/-^ adipose tissue exhibited an increased basal oxygen consumption rate (OCR; Figure 3F). Accordingly, the levels of pyruvate dehydrogenase kinase 4 (PDK4), which inhibits pyruvate dehydrogenase (PDH) complex, thereby shifting nutrient flux towards fatty acid oxidation, was also higher in *vdr*^-/-^ adipose compared to WT mice (Figure 3G).

**Fig. 3.**
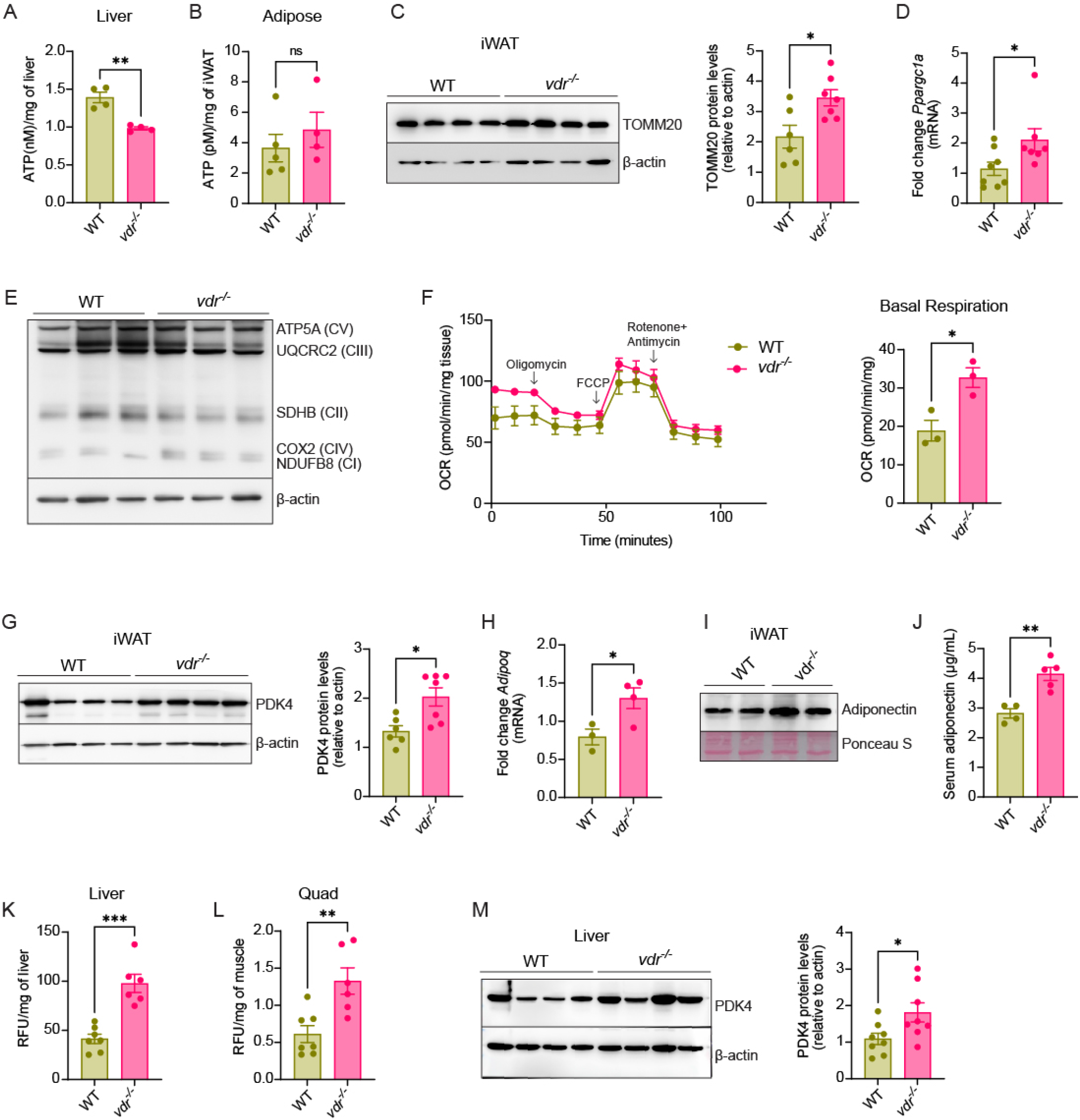
*vdr* ^-/-^ mice show increased basal mitochondrial activity in the adipose. (A&B) ATP levels estimated in the liver (A) and iWAT (B) of 7-week WT and *vdr* ^-/-^ mice. (C) Representative western blot and quantification (right panel) for TOMM20 in iWAT of 7-week WT and *vdr* ^-/-^ mice. (D) mRNA expression levels of *Ppargc1a* in iWAT of 7-week WT and *vdr* ^-/-^ mice as quantified using qRT-PCR. (E) A representative western blot using a cocktail of antibodies against mitochondrial electron transport chain components in iWAT of 7-week WT and *vdr* ^-/-^ mice. (F) Seahorse flux analysis depicting the OXPHOS activity and basal respiration measurement in iWAT explants of 7-week WT and *vdr* ^-/-^ mice. The time of addition of various inhibitors is shown by arrows. (G) Representative western blot and quantification (right panel) of PDK4 in iWAT of 7-week WT and *vdr* ^-/-^ mice. (H) Expression levels of adiponectin mRNA in iWAT of 7-week WT and *vdr* ^-/-^ mice as quantified using qRT-PCR. (I) Western blot showing the protein levels of adiponectin in iWAT of 7-week WT and *vdr* ^-/-^ mice. Ponceau stained blot is given to show equal loading. (J) Serum levels of adiponectin in 7-week WT and *vdr* ^-/-^ mice, measured using ELISA. (K&L) Bodipy-C12 uptake in liver (K) and Quad muscle (L) of 7-week WT and *vdr* ^-/-^ mice. (M) Representative immunoblot and quantification (right panel) of PDK4 in liver of 7-week WT and *vdr* ^-/-^ mice. All graphs show mean ± SEM. Statistical significance was determined by unpaired t-test (*p < 0.05, **p < 0.01, ***p< and ****p < 0.0001). The number of samples is denoted by the dots in the graphs.

Conditions that lead to energy deprivation induce adipose tissue to secrete the adipokine adiponectin, which modifies the metabolic behaviour of tissues and enhances mitochondrial function in adipocytes (42). Interestingly, the mRNA and protein levels of adiponectin were high in *vdr*^-/-^ adipose as compared to WT (Figure 3H&I), which is in line with studies showing an inverse correlation of adiponectin levels with adipose tissue mass (43). The serum adiponectin levels were also higher in *vdr*^-/-^ mice at 7 weeks of age than WT (Figure 3J). Intriguingly, the changes in serum levels of adiponectin at 3, 5, or 6 weeks of age were comparable between WT and *vdr*^-/-^ mice (Figure S3B), implying that adiponectin upregulation occurs as a result of progressive energy deprivation and adipose atrophy. Apart from adipose, adiponectin production has been observed in several other tissues, including muscle (44). We observed higher adiponectin levels in the quadricep muscle of *vdr*^-/-^ as compared to WT at 7 weeks of age (Figure S3C). Adiponectin is shown to increase fatty acid uptake & oxidation by skeletal muscle and liver (45, 46). Accordingly, we observed higher fluorescence intensity of BODIPY-C12 in the liver and skeletal muscle (Quad) of *vdr*^-/-^, suggesting increased lipid uptake by these tissues (Figure 3K&L). Further, the liver of *vdr*^-/-^ also exhibits increased PDK4 expression (Figure 3M). All these results suggest an overall shift towards fatty acid oxidation in *vdr*^-/-^ adipose and liver.

### *vdr*-/-mice show defective liver glycogenolysis and increased protein phosphatase 1 level

Taken together, we argue that the adipose wasting observed in the *vdr*^-/-^ mice is caused by systemic energy deprivation. Muscle glycogen storage defect previously observed in these mice could be one of the underlying factors that cause energy deficiency (14, 18). However, it does not fully explain the reduced blood glucose levels in these mice. During fasting, the liver breaks down glycogen stores via glycogenolysis and initiates gluconeogenesis to produce glucose and maintain circulating blood glucose levels (47). To address whether the inability of *vdr*^-/-^ mice to maintain plasma glucose concentrations is due to defects in gluconeogenesis, we performed a pyruvate tolerance test. WT and *vdr*^-/-^ mice were fasted for 12 hours followed by an intraperitoneal injection of pyruvate. Measurement of glucose level post pyruvate injection at different time points revealed that total hepatic glucose production was not altered in *vdr*^-/-^ mice (Figure 4A). Gluconeogenesis-related genes, phosphoenolpyruvate carboxykinase (*Pepck*), and fructose-1,6-bisphosphatase (*Fbp1*) were also unaffected in the livers of WT and *vdr*^-/-^ mice (Figure S4A&B), indicating that gluconeogenesis is not affected in these mice.

**Fig. 4.**
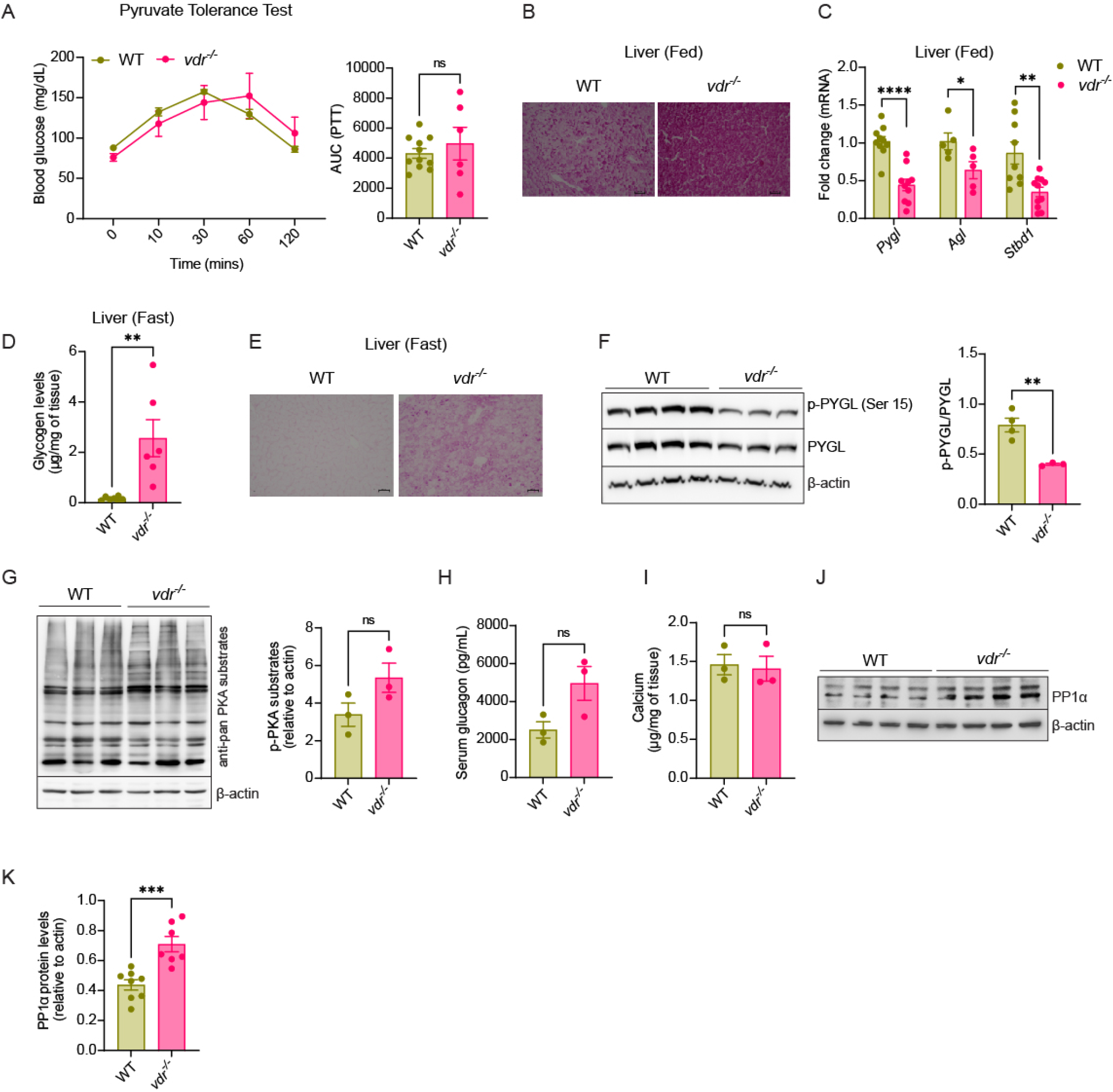
*vdr* ^-/-^ mice show defective liver glycogenolysis and increased protein phosphatase 1 level. (A) Pyruvate tolerance test and Area Under the Curve (AUC) in 7-week WT and *vdr* ^-/-^ mice. (B) Micrographs of PAS-stained liver sections (*ad libitum* fed state) in 7-week WT and *vdr* ^-/-^ mice. (C) mRNA expression levels of glycogen phosphorylase (*Pygl*), glycogen debranching enzyme (*Agl*), and starch binding domain-containing protein 1 (*Stbd1*) in the liver of 7-week WT and *vdr* ^-/-^ mice as quantified using qRT-PCR under *ad libitum* fed state. (D) Glycogen content (µg/mg of tissue) in 12 hour-fasted liver in 7-week WT and *vdr* ^-/-^ mice. (E) Micrographs of PAS-stained liver sections (12 hour-fasted) in 7-week WT and *vdr* ^-/-^ mice. (F) Western blot representing the levels of phospho-PYGL (Ser 15), PYGL, and β-actin and the ratio of phospho-PYGL to total PYGL (right panel) in the liver (12 hour-fasted) of 7-week WT and *vdr* ^-/-^ mice. (G) Representative western blot and quantification (right panel) for phospho-PKA substrates (RRXS*/T*) in the liver (12 hour-fasted) of 7-week WT and *vdr* ^-/-^ mice. (H) Serum glucagon levels in 7-week WT and *vdr* ^-/-^ mice after 12 hours of fasting. (I) Calcium levels (µg/mg of tissue) in the liver of 7-week WT and *vdr* ^-/-^ mice after 12 hours of fasting measured via ICP-MS. (J&K) Representative western blot (J) and quantification (K) for PP1α in the fasted liver of 7-week WT and *vdr* ^-/-^ mice. All graphs show mean ± SEM. Statistical significance was determined by unpaired t-test (*p < 0.05, **p < 0.01, ***p< 0.001 and ****p < 0.0001). The number of samples is denoted by the dots in the graphs.

Liver glycogenolysis provides a major source of fuel, which is mobilized during the early phase of starvation (20). Intriguingly, and contrary to expectations, we observed that *vdr*^-/-^ mice displayed increased glycogen levels as measured by periodic acid Schiff (PAS) staining under fed conditions (Figure 4B). mRNA levels of glycogenin, an enzyme involved in the initial step of glycogen synthesis, were unaffected in *vdr*^-/-^ mice (Figure S4C). There were no differences in mRNA and protein levels of glycogen synthase (Figure S4D&E). However, mRNA levels of glycogen phosphorylase (*Pygl*), glycogen debranching enzyme (*Agl*), and starch binding domain 1 (*Stbd1*) protein involved in glycogen degradation were reduced in *vdr*^-/-^ as compared with WT, suggesting a derangement in the glycogen degradation machinery (Figure 4C). The reduction in glycogen degrading enzymes was not observed in 3 weeks *vdr*^-/-^ mice (Figure S4F). To confirm a glycogen degradation defect, we analyzed the liver glycogen levels after fasting the mice for 12 hours. *vdr*^-/-^ mice exhibited a drastic decrease in glycogen utilization, suggested by higher glycogen levels by biochemical estimation and PAS staining (Figure 4D&E). To decipher the molecular mechanisms underlying reduced glycogenolysis during fasting in *vdr*^-/-^ mice, we examined the activity of glycogen phosphorylase (PYGL), one of the key enzymes in glycogen degradation pathway. Western blot analysis of fasted liver samples from *vdr*^-/-^ showed reduced PYGL activity, as suggested by a profound reduction in the activating Ser15 phosphorylation (Figure 4F).

Next, we sought to delineate the mechanism underlying reduced glycogen phosphorylase activity in *vdr*^-/-^. The activity of glycogen phosphorylase is regulated via the phosphorylation-dephosphorylation cycle by glycogen phosphorylase kinase and protein phosphatase 1, respectively. We asked if the reduced activity of glycogen phosphorylase kinase, which phosphorylates PYGL, accounts for low PYGL activity. To address this, we measured the activity of PKA, a protein kinase that phosphorylates glycogen phosphorylase kinase, in the liver of WT and *vdr*^-/-^ mice. Immunoblotting analysis using antibodies specific for protein kinase A (PKA) substrates revealed that phosphorylation of most of the PKA substrates was similar in the liver of WT and *vdr*^-/-^ mice (Figure 4G). Moreover, fasting levels of circulating glucagon, which is one of the major drivers of hepatic glycogenolysis and work via PKA activation, remain unchanged in the *vdr*^-/-^ mice as compared to WT (Figure 4H), confirming similar activation of the PKA-related signaling pathway in WT and *vdr*^-/-^ liver. Glycogen phosphorylase kinase activity is also regulated by calcium dynamics, and *vdr*^-/-^ mice exhibit reduced serum calcium levels at 7 weeks (Figure S4G). So, we checked if differences in intracellular calcium levels are linked to reduced glycogen phosphorylase activity. ICP-MS analysis did not find any statistically significant difference in the intracellular calcium levels between the livers of *vdr*^-/-^ and WT mice (Figure 4I). Calcium-dependent signalling is also unaffected, as suggested by similar activation of calcium/calmodulin dependent protein kinase II (CaMKII) in WT and *vdr*^-/-^ liver (Figure S4H). We then measured the protein levels of the catalytic (PPP1cα) subunit of protein phosphatase 1 (PP1), which inactivates PYGL by dephosphorylation. Compared to WT mice, *vdr*^-/-^ mice show a significant increase in PPP1cα level (Figure 4J&K). Altogether, these results show that the liver of *vdr*^-/-^ mice exhibits a glycogen utilization defect characterized by decreased PYGL activity and increased protein phosphatase 1 levels.

### Weaning on a calcium-rich diet prevents adipose wasting and rescues glycogenolysis defect in *vdr*^**-/-**^

*vdr*^-/-^ mice are indistinguishable from WT littermates till the weaning stage (11, 12). Notably, 2 weeks post-weaning, *vdr*^-/-^ mice fed a regular chow diet show abnormalities such as hypocalcemia, and growth retardation, and ultimately die within 4 weeks post-weaning. Levels of serum calcium at 3 weeks of age (weaning) are similar between WT and *vdr*^-/-^ (Figure S5A), which is significantly reduced post-weaning (Figure S4G). To elucidate if VDR elicits its effect on carbohydrate utilization through calcium regulation, we maintained the normocalcemic status of *vdr*^-/-^ mice by weaning them onto a high-calcium rescue diet (48). Serum calcium levels at 7 weeks of age were similar between *vdr*^-/-^ on the rescue diet and WT mice (Figure S5B). Reduced serum calcium in *vdr*^-/-^ is associated with elevated PTH level (16), which was reduced when they were subjected to the rescue diet (Figure S5C). Furthermore, the rescue diet restored body weight (Figure 5A) and muscle weight (TA, Quad, and Gas) in *vdr*^-/-^ mice (Figure S5D). In consonance with restoration of muscle weight, grip strength of *vdr*^-/-^ on the rescue diet was similar to that of WT (Figure S5E).

**Fig. 5.**
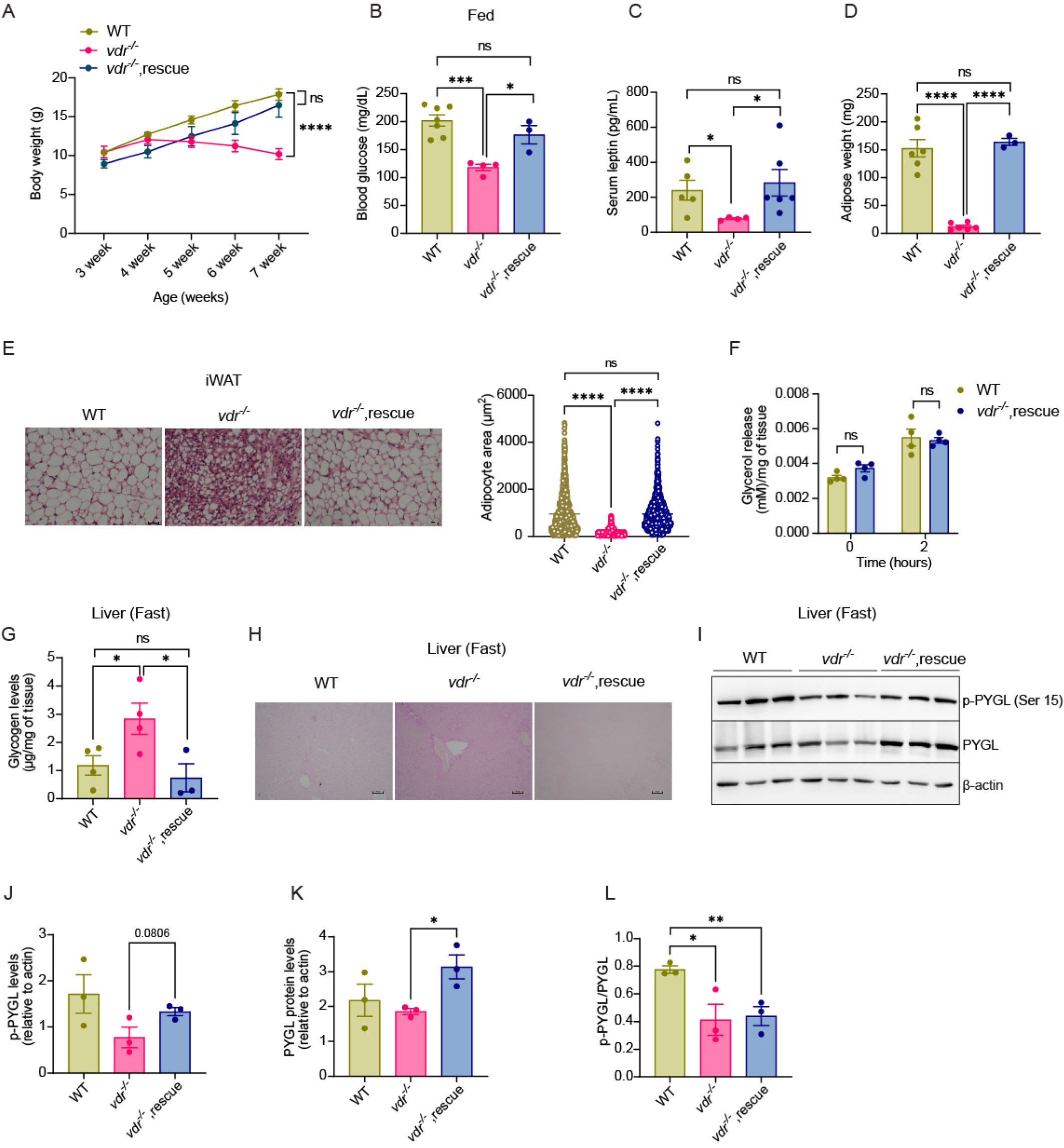
Weaning on a calcium-rich diet prevents adipose wasting and rescues glycogenolysis defect in *vdr* ^-/-^. (A) Body weight (in grams) of WT and *vdr* ^-/-^ mice on chow diet and *vdr* ^-/-^ on rescue diet from 3 to 7-weeks (B) Blood glucose levels (in mg/dL) in 7-week WT and *vdr* ^-/-^ mice on chow diet and *vdr* ^-/-^ on rescue diet under *ad libitum* fed state. (C) Serum levels of leptin (in pg/mL) in 7-week WT and *vdr* ^-/-^ mice on chow and rescue diet measured using ELISA under *ad libitum* fed state. (D) Inguinal WAT weight (in mg) of 7-week WT and *vdr* ^-/-^ mice on chow and rescue diet. (E) Representative images of H&E stained iWAT sections and quantitative analysis of adipocyte area in 7-week WT and *vdr* ^-/-^ mice on chow and rescue diet. (F) Glycerol release from WAT explants of 7-week WT and *vdr* ^-/-^ mice on chow and rescue diet. (G) Glycogen content (µg/mg of tissue) in the fasted liver of 7-week WT and *vdr* ^-/-^ mice on chow and rescue diet. (H) Micrographs of PAS-stained fasted liver sections in 7-week WT and *vdr* ^-/-^ mice on chow and rescue diet. (I) Western blot representing the levels of phospho-PYGL (Ser 15), PYGL, and β-actin. (J-L) Quantification of phospho-PYGL relative to actin (J), total-PYGL relative to actin (K), and phospho-PYGL relative to total-PYGL (L) in fasted liver of 7-week WT and *vdr* ^-/-^ mice on chow and rescue diet. All graphs show mean ± SEM. Statistical significance was determined by unpaired t-test (*p < 0.05, **p < 0.01, ***p< 0.001 and ****p < 0.0001). The number of samples is denoted by the dots in the graphs.

Interestingly, high-calcium diet supplementation effectively abolished energy insufficiency in *vdr*^-/-^ mice as blood glucose levels and leptin levels were maintained to WT levels in *vdr*^-/-^ on rescue diet (Figure 5B&C). These mice did not show adipose wasting as adipose tissue weight was comparable between WT and *vdr*^-/-^ on rescue diet (Figure 5D). Histopatho-logical analysis of adipose sections confirmed this (Figure 5E). *vdr*^-/-^ mice maintained on the rescue diet display similar levels of adipose lipolysis as that of the WT mice, further supporting the restoration of energy homeostasis in these mice (Figure 5F). Furthermore, hepatic glycogen levels in *vdr*^-/-^ on the rescue diet were significantly lower as compared to *vdr*^-/-^ mice on regular chow, clearly showing that liver glycogen utilization is restored by the rescue diet (Figure 5G). PAS staining substantiates reduced accumulation of glycogen in the *vdr*^-/-^ rescue group as compared to the *vdr*^-/-^ mice on regular chow diet (Figure 5H). Next, we checked if the PYGL regulation is restored in these mice (Figure 5I). Interestingly, while there was a general increase in the Ser15 phosphorylated PYGL (calculated against actin), it was primarily due to an increase in the PYGL protein level rather than the phosphorylation as evidenced by no change in the ratio of phosphorylated versus total protein levels (Figure 5I-L). However, this indicates that there is a larger pool of active PYGL, which aligns with the improved utilization of glycogen. Ablation of the vitamin D signaling in global *vdr*^-/-^ mice leads to impaired serum insulin levels, which was corrected by a rescue diet in *vdr*^-/-^ mice (Figure S5F). Overall, these results indicate that normalizing serum calcium levels in *vdr*^-/-^ mice could restore energy homeostasis, prevent hepatic glycogenolysis defect and effectively abolish adipose wasting.

### Milk-fat diet supplementation circumvents the metabolic defects induced by VDR deficiency without normalizing serum calcium levels

Data shown so far indicates that the VDR-calcium axis is important for the metabolic transition during weaning. If this is true, we hypothesized that the mice continued on a high-fat-containing milk-based diet, should maintain energy homeostasis. We tested this by weaning these mice to a custom-designed milk-based diet (MFD) beginning at day 21 and continuing through adulthood till 7 weeks. The body weight of *vdr*^-/-^ mice on MFD was comparable to WT litter mates (Figure 6A). No mortality was recorded during the period of the study. *vdr*^-/-^ mice on MFD showed blood glucose levels similar to that of WT on chow, indicating that MFD alleviated the energy deficiency in *vdr*^-/-^ mice (Figure 6B). Concordantly, the adipose weight was also preserved in these mice when subjected to MFD (Figure 6C). Furthermore, the adipose lipolysis was reduced to that of WT levels in MFD-fed *vdr*^-/-^ mice (Figure 6D). Histopathological analysis of adipose sections also indicated similar adipose area between WT & *vdr*^-/-^ MFD (Figure 6E), clearly indicating that the adipose atrophy is alleviated in these mice. The ability to restore adipose tissue is also supported by our previous data showing the adipogenic and lipogenic gene expression was unchanged in the adipoe tissue of these mice (Figure 1). Rather, the adipose atrophy is a response to systemic energy deprivation. Moreover, glycogen metabolism in the liver was also restored in *vdr*^-/-^ mice on MFD as confirmed by higher glycogen content after 12 hour fasting using biochemical estimation (Figure 6F) and PAS staining (Figure 6G). Further, the restoration glycogen metabolism in MFD fed mice was confirmed by increased levels of PYGL protein (Figure 6H&I). Prevention of the onset of metabolic defects by MFD was not dependent upon normalisation of serum calcium levels as calcium levels in *vdr*^-/-^ on MFD was similar to *vdr*^-/-^ chow (Figure 6J). Thus, altogether, these experimental findings demonstrate that a milk-based diet can ameliorate the defect in hepatic glycogen utilization in *vdr*^-/-^ by some alternate pathways that do not depend on the normalization of serum calcium.

**Fig. 6.**
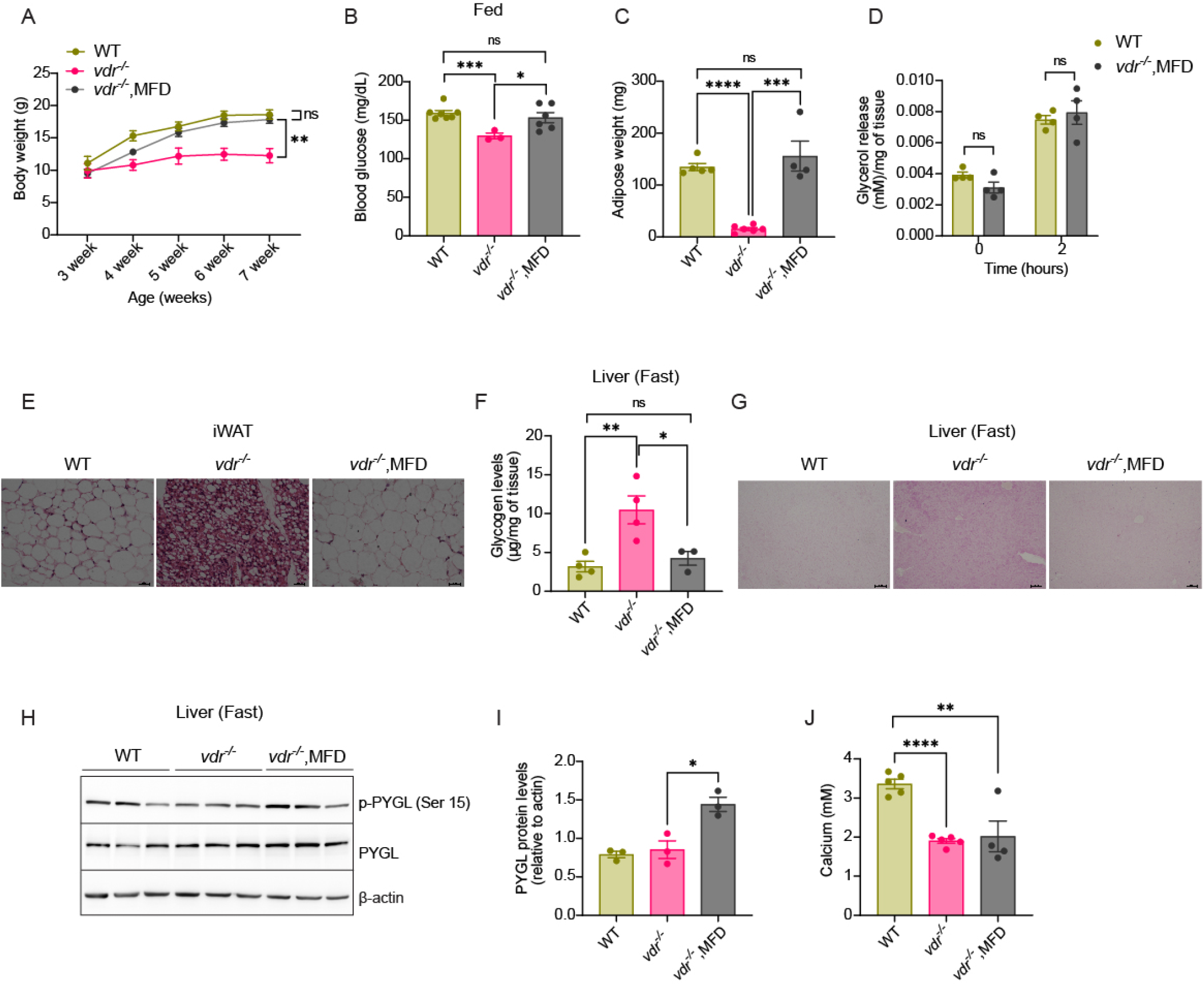
Milk-fat diet supplementation circumvents the metabolic defects induced by VDR deficiency without normalizing serum calcium levels. (A) Body weight (in grams) of WT and *vdr* ^-/-^ mice on chow and MFD from 3 to 7 weeks. (B) Blood glucose levels (in mg/dL) in 7-week WT and *vdr* ^-/-^ mice on chow diet and MFD. (C) Inguinal WAT weight (in mg) in 7-week WT and *vdr* ^-/-^ mice on chow diet and MFD. (D) Glycerol release from iWAT explants of 7-week WT and *vdr* ^-/-^ mice on chow diet and MFD. (E) Representative images of H&E stained iWAT sections in 7-week WT and *vdr* ^-/-^ mice on chow and MFD. (F) Glycogen content (µg/mg of tissue) in the fasted liver in 7-week WT and *vdr* ^-/-^ mice on chow diet and MFD. (G) Micrographs of PAS-stained fasted liver sections in 7-week WT and *vdr* ^-/-^ mice on chow and MFD diet. (H&I) Western blot representing the levels of phospho-PYGL (Ser 15), total PYGL and β-actin (H) and accompanying quantification of total PYGL relative to actin (I) in the fasted liver of 7-week WT and *vdr* ^-/-^ mice on chow diet and MFD. (J) Serum calcium levels (in mM) of 7-week WT and *vdr* ^-/-^ mice on chow and MFD diet, measured using ELISA. All graphs show mean ± SEM. Statistical significance was determined by unpaired t-test (*p < 0.05, **p < 0.01, ***p< 0.001 and ****p < 0.0001). The number of samples is denoted by the dots in the graphs.

## Discussion

During the weaning process, the systemic metabolic machinery must adapt to utilize and store the higher amount of glucose in the diet. However, the factors that enable this transition are not known. The data presented here show that vitamin D receptor signaling is essential for the metabolic shift during the weaning process. *vdr*^-/-^ mice exhibit severe energy deprivation after weaning, characterized by severe adipose wasting seemingly driven by increased norepinephrine levels in *vdr*^-/-^ adipose. Importantly, the wasting phenotype is alleviated by restoration of normocalcemia in these mice by providing a high-calcium rescue diet, indicating that the VDR-calcium axis is essential for metabolic adaptation during weaning. Indeed, the ability of the milk-fat diet to prevent energy deprivation and wasting in these mice supports the hypothesis that the VDR-calcium axis is essential for the adaptation of animals to carbohydrate-based diets during weaning.

A significant feature of *vdr*^-/-^ when weaned onto a carbohydrate-based diet is the defective glycogen handling in the liver and muscles (18). We earlier showed that *vdr*^-/-^ mice fail to secrete insulin in response to glucose stimulation (14). The hypoglycemia exhibited by *vdr*^-/-^ despite lower serum insulin levels was perplexing. In this study, we demonstrate that *vdr*^-/-^ mice fail to efficiently utilize stored hepatic glycogen due to low activity of the catabolic enzyme PYGL. Notably, patients affected by Hers disease, an inherited glycogen storage disease caused by deficiency of PYGL, present hypoglycemia, and growth retardation (49) as seen in *vdr*^-/-^ mice. These patients also show elevated triglyceride levels, which are considered a manifestation of increased lipolysis (50), which resembles the liver-adipose defects observed in the *vdr*^-/-^ mice.

Interestingly, though the skeletal muscles of *vdr*^-/-^ also show a glycogen storage defect, it appears to be caused by a different mechanism. In the muscles, glycogen accumulation is associated with increased glycogen synthase protein levels and reduced phosphorylation, presumably due to lower PKA activity. This suggested that increased glycogen synthesis plays an important role in the glycogen accumulation in the muscles of *vdr*^-/-^ mice (18). However, in the liver, we did not find any change in the glycogen synthase protein levels or phosphorylation. On the other hand, glycogen accumulation in the liver is driven by reduced activity of glycogen phosphorylase. Moreover, glucagon levels and PKA activities were unchanged in the liver. On the other hand, protein phosphatase 1 levels were significantly high in the *vdr*^-/-^ liver. Because calcium is a well-known activator of glycogenolysis, it is possible that the lower calcium availability could also be one of the reasons for the abnormal glycogen accumulation in *vdr*^-/-^ liver. Albeit serum calcium levels are low in *vdr*^-/-^, intracellular hepatic calcium levels and calcium signaling that regulate PYGL are intact, making it unlikely that the observed dysfunctional glycogenolysis is attributable to changes in calcium. Still, the indirect effect of calcium on neural outputs to peripheral organs through the brain-liver axis cannot be entirely excluded.

Starvation and other energy deprivation conditions can induce adipocyte lipolysis through the sympathetic nervous system (51). Defects in glycogen breakdown from hepatic stores, combined with skeletal muscle glycogen storage disorder (18), may induce energy deprivation and stimulate adipose tissue lipolysis. Indeed, the increased levels of adipose tissue norepinephrine and lipolysis support this hypothesis. Furthermore, the loss of adipose mass in *vdr*^-/-^ could neither be attributed to changes in the expression of genes associated with lipogenesis and adipogenesis nor to increased thermogenesis. Apart from the systemic negative energy balance, other factors are also known to affect adipose mass and lipolysis. Previous studies have demonstrated adipose lipolysis to be inversely correlated with expression levels of VDR (52, 53). However, whether the increased lipolysis in adipose we observed is driven by lack of VDR expression in the adipocytes is not apparent. Furthermore, inflammatory cell infiltration in the adipocytes is also known to induce adipose atrophy (54). However, we did not observe any evidence of increased inflammation in *vdr*^-/-^ adipose. Taken together, the data strongly indicate that the lipolysis in *vdr*^-/-^ is caused by systemic energy deprivation and the resultant increase in the norepinephrine signaling.

Mice with systemic genetic ablation of VDR are phenotypically normal till weaning but exhibit symptoms of rickets/osteomalacia and secondary hyperparathyroidism primarily after weaning (11). We and others have shown normal calcium levels in *vdr*^-/-^ mice at 3 weeks, which drops down significantly at 7 weeks (17). Our data show that lipodystrophy and defective hepatic glycogenolysis are not observed in 3-week-old mice, whereas 7-week-old *vdr*^-/-^ mice develop metabolic defects, including adipose wasting, reduced blood glucose and leptin levels, indicative of a starvation-like state. This suggest that lack of VDR signalling impairs metabolic adaptation during weaning, decreasing the ability to switch off fatty acid oxidation during the transition process. Restoration of circulating calcium concentrations in *vdr*^-/-^ by maintaining them onto a “rescue diet,” after weaning ameliorates negative energy balance, abolishes adipose wasting and rescues glycogenolysis defects. In support of this, calcium infusions have been found useful in patients with hereditary resistance to 1,25(OH)2D to correct biochemical and bone abnormalities (55).

When we allowed *vdr*^-/-^ mice to fed on a diet enriched in fat from day 21 and continuing through adulthood, the defects in adipose mass and glycogenolysis were alleviated without restoring calcium homeostasis. This clearly shows that even in the absence of the VDR-calcium axis, an alternate fat-based metabolic state can be established in these mice. This emphasizes the crucial role of VDR in supporting efficient metabolic adaptation during the transition from a calcium-rich milk-based diet to a carbohydrate diet after weaning. Consolidating the observations made here and in our previous studies, we found at least three post-weaning defects in *vdr*^-/-^ mice that can directly affect glucose homeostasis: Glycogenolysis defects in the liver as reported in this study, abnormal glycogen storage in the skeletal muscles (18), and defective insulin response (14). All three of these defects were alleviated by containing these mice on a milk-fat-containing diet. Clearly, further investigations are required to discern the origin of the molecular pathways that initiate the metabolic derangement in the absence of VDR, and which of these factors plays the key etiological role needs to be addressed.

## Materials and Methods

### Animal maintenance and Diet

VDR null mutant mice were purchased from the Jackson Laboratory (Stock No. 006133, B6.129S4-Vdrtm1Mbd lJ; Bar Harbor, ME, USA) and housed in sterile, individually ventilated cages. All animals were maintained in a temperature-controlled environment with a 12 hour-light/dark cycle and were weaned onto commercial rodent chow (Altromin, 1314), Milk Fed Diet (Research Diets, D19112203), or calcium-rich rescue diet (20% lactose, 2% calcium and 1.25% phosphorus, VRK Nutritional Solutions). Mice were euthanized at 3 or 7 weeks of age, and samples were collected in a fasted (12-hour) or fed (*ad libitum*) state and frozen at -80°C until use. Maintenance of animal strain, tissue collection after euthanasia and drug administration were performed as per the guidelines of the animal ethics committee of the National Institute of Immunology.

### Western blotting

Frozen adipose and liver tissues were homogenized in lysis buffer (50 mM Tris–HCl pH 7.5, 150 mM NaCl, 5 mM EDTA, 1% NP-40, 0.5% sodium deoxycholate, 0.1% SDS) supplemented with protease/phosphatase inhibitor cocktail (5872, Cell signalling). Homogenates were centrifuged at 13000 rpm for 15 minutes at 4°C. The supernatant was collected, and protein quantification was done using the BCA method. An equal amount of protein for each sample was resolved by SDS-PAGE (8%, 10%, 12%, and 15%) and transferred onto a nitrocellulose membrane (Amersham). The membrane was incubated with primary antibodies, followed by incubation with HRP-conjugated anti-mouse or anti-rabbit secondary antibodies, and the protein bands were visualized using a chemiluminescent substrate (Bio-Rad, 1705060). Chemiluminescence was acquired with the LAS 500 Imager system (GE). β-actin was used as a loading control. Quantification of western blots was performed using ImageJ software. List of antibodies is given in Table S1.

### RNA extraction and quantitative real-time PCR

Adipose tissue and liver were dissected and snap-frozen in liquid nitrogen and homogenized in RNAiso plus Reagent (Takara, 1 ml/100 mg tissue) to isolate total adipose and liver RNA as per the manufacturer’s recommendations. Total RNA was quantified with a nanodrop 8000 spectrophotometer (Thermo Scientific, Wilmington), and a ratio of 2 for the absorbance of 260 to 280 nm was used to determine the RNA quality. Complementary DNA (cDNA) was then synthesized from 1 µg of RNA using a reverse transcription kit (6110A; Takara). Quantitative RT–PCR was performed using the QuantStudio-6 Flex Real-Time PCR Systems (Thermo Fisher Scientific) with SYBR® Premix Ex Taq (Tli RNase H Plus, RR420A; Takara). Each sample was amplified in triplicates using gene-specific primers. The expression levels of each transcript were normalized to the housekeeping gene L7. The expression of each target gene was calculated as the ‘fold change’ relative to the control samples according to the 2-L}L}Ct method. The primer sequences used are mentioned in Table S2.

### ATP estimation in adipose tissue and liver

ATP estimation was done using a bioluminescence-based assay kit (A22066; Thermo Scientific). Briefly, Tissues were chopped into small pieces and sonicated in a modified RIPA buffer with EDTA (150 mM NaCl, 1% NP40, 0.5% Na-deoxycholate, 0.1% SDS, 25 mM Tris, pH 7.4) and centrifuged at 13000 rpm for 15 min at 4°C. The supernatant was collected and proceeded with the assay according to the manufacturer’s protocol. Levels of ATP were normalized to tissue weight.

### Histological analysis

Adipose tissue and liver were dissected out, weighed, fixed in 10% formalin (wt/vol.), and embedded in paraffin. Histological sections (7 µm) of inguinal white adipose tissues and liver were mounted on glass slide and stained with hematoxylin and eosin (H&E) and periodic acid-Schiff (PAS) according to the manufacturer’s protocol (Sigma). Five digital images from non-overlapping fields were taken of each slide (n>3). ImageJ software (National Institutes of Health, Bethesda, MD, USA) was used to quantify the cross-sectional area of adipocytes in 3–4 fields per slide.

### Ex-vivo lipolysis assay

Freshly dissected inguinal white adipose tissue was minced into small pieces and incubated in Krebs-Ringer bicarbonate-HEPES buffer supplemented with 3% fatty acid free BSA and 5 mM glucose at 37 °C for 2 h with gentle shaking. Following the incubation, aliquots were collected and assayed for glycerol content using glycerol determination kit (MAK117, Sigma) according to the manufacturer’s instruction. Glycerol levels were normalized to tissue weight.

### Pyruvate tolerance test

WT and homozygous VDR mutant mice were fasted for a period of 12 hours. During that period, the mice had free access to water. Sodium pyruvate (1 g/kg body weight) was administered at time 0 by intraperitoneal injection in saline. Blood glucose levels at 0, 10, 30, 60, and 120 minutes were determined using blood from the tail vein using the Accu-Check Instant glucometer.

### Estimation of adiponectin levels in serum and muscle

Serum adiponectin levels were estimated using mouse ADP/Acrp30(Adiponectin) ELISA kit (E-EL-M0002) according to the manufacturer’s instructions. Muscle adiponectin levels were estimated using muscle lysates prepared in the assay buffer as per the manufacturer’s protocol. The protein level of muscle lysates was quantified by the BCA method. Levels of muscle adiponectin were normalized to the muscle protein levels.

### Estimation of hormone levels in serum and tissue

Serum adrenaline levels were estimated using an adrenaline ELISA kit (ab287788) according to the manufacturer’s protocol. Serum PTH levels were estimated using GENLISA™ mouse parathyroid hormone (PTH) ELISA (KLM0560) according to the manufacturer’s protocol. Serum insulin levels were estimated in *ad libitum*-fed mice using a rat/mouse insulin ELISA kit (EZRMI-13K) according to the manufacturer’s protocol. Serum levels of leptin, ghrelin, and glucagon were estimated using Bio-Plex Pro mouse diabetes 8-plex Assay (171F7001M) as per the manufacturer’s protocol. Serum and iWAT norepinephrine levels were estimated using norepinephrine ELISA kit (ab287789) according to the manufacturer’s instructions. Levels of norepinephrine in iWAT were normalized to tissue weight.

### Estimation of glycogen content in liver tissue

50 mg of liver tissue was taken and glycogen was quantified in fasted state using Glycogen assay kit (ab65620) according to the manufacturer’s protocol. Levels of glycogen in liver was normalized to tissue weight.

### Intracellular calcium estimation using ICP-MS (Inductively coupled plasma mass spectrometry)

50 mg of liver tissue was taken in a digestion vial (64MG5 vials) and incubated with 70% HNO3 (400 µl) and H2O2 (100 µl) for 20 min at room temperature. Vials were then sealed and subjected to microwave digestion. The initial power was set at a ramp of 150 W for 10 min with a max temperature of 140°C, followed by a 15 min hold. Then, a second ramp of 250 W for 10 min was used, and a final hold was used to bring the temperature to 55°C at the highest fan speed. After diluting the samples with trace metal-free water, calcium levels were monitored in the samples using ICP-MS (iCAPTM TQ ICP-MS, Thermo Scientific, USA). Thermo Scientific QtegraTM Intelligent Scientific Data Solution (ISDS) software was used to operate and control the instrument. Briefly, the torch was warmed up for 30 min in single-quad kinetic energy discrimination (SQ-KED) mode with helium to remove the polyatomic nuclei interference and then autotuned in normal mode followed by advanced KED mode. Then, the multi-element standard (92091, Sigma Aldrich, USA) of different concentrations was run at the same settings to prepare the standard plot. A sample blank was run before each sample, and a quality control (QC) of known concentrations of standards was run after every 15–20 samples to monitor and maintain uniform signal intensity throughout the run. Data analysis was done using Qtegra software (Thermo Scientific, USA).

### Serum calcium estimation

Serum calcium levels were determined using calcium estimation kit (ab102505) according to manufacturer instructions.

### Bodipy uptake assay

For fatty acid uptake assay, mice were given an intraperitoneal injection of 40 µM Bodipy C12 (3822, Thermo Fisher Scientific) dissolved in 100 µl of 1X PBS. After 1.5 hours of Bodipy administration, liver and muscle samples were collected and were homogenized in the RIPA buffer. The supernatant was collected, and fluorescence was measured (excitation-488nm, emission-520nm). Fluorescence was normalized to tissue weight.

### Flow cytometry analysis of adipose tissue

Inguinal WAT was excised and minced in a digestion buffer containing 1X HBSS, 3% BSA, 1 mg/mL collagenase 1 (TC211), 1.2 mM CaCl2, and 1 mM MgCl2. The minced tissue was incubated at 37°C for 1.5 hours with shaking. The suspension was passed through a 70 µm strainer to remove the undigested clumps and debris. The suspension was centrifuged at 400 g for 10 mins at 4°C. The supernatant containing floating adipocytes was discarded, and the stromal vascular fraction (SVF) pellet was resuspended in 1X PBS. For FACS staining, cells were washed with 1X PBS and resuspended in staining buffer (1X PBS+ 2% FBS) containing Fc blocker (1:1000), and the following conjugated antibodies: CD45-APC-Cy7 (1:400), CD11b-FITC (1:200), CD11c-PE (1:200), F4/80-BV421 (1:200), CD206-APC (1:400) and viability dye, eflour 506 (1:1000). The cells were incubated at 4°C for 30 mins followed by washing with PBS. Samples were analyzed by flow cytometry (BD LSRFortessa X-20).

### Grip strength

Muscle strength was evaluated in different groups using a grip strength meter (GT3, Panlab, Harvard Apparatus). Mice were lifted by the base of the tail and placed so that their paws gripped the trapeze. Each mouse was tested five times with a short rest between each set. An average of three tests was recorded. Results are expressed as gram force.

### Extracellular metabolic flux analysis

Adipose tissue oxygen consumption rate was assessed using the seahorse XFe24 analyzer (Agilent Technologies, Santa Clara, CA, USA) according to the manufacturer’s specifications. Briefly, the seahorse cartridge was hydrated overnight in a non-CO2 incubator. On the day of the assay, freshly isolated inguinal white adipose tissue (5 mg) was rinsed with 1X PBS and cut into pieces (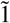
 mg), washed extensively, and then each piece was placed in a single well of an XF24-well Islet Flux plate (#103518; Agilent Technologies, Santa Clara, CA, USA). The tissue was covered with a customized screen, which allowed free perfusion while minimizing tissue movement. XF basal medium supplemented with 10 mM glucose, 1 mM pyruvate, and 2 mM glutamine was then added to each well. The plate was incubated at 37°C in a non-CO2 incubator for 45 minutes before extracellular flux analysis. The mitochondrial stress test was performed using subsequent injections of 100 µM oligomycin (75351, Sigma-Aldrich), 100 µM FCCP (C2920, Sigma-Aldrich), 60 µM rotenone (R8875, Sigma-Aldrich)/antimycin (A8674, Sigma-Aldrich). Data was analyzed using the seahorse wave software v2.6.3.5 (Agilent Technologies). Oxygen consumption rate values were normalized to tissue weight.

### Statistical analysis

Statistical analysis was performed using the GraphPad software (Prism 9) using unpaired student’s two-tailed t-tests, (*p<0.05, **p<0.01, ***p<0.001,****p<0.0001, ns=not significant). This is also specified in the figure legends. Numerical values for each measurement are displayed as mean±standard error mean (SEM).

## Ethical Standards Statement

The authors of this manuscript certify that all experiments in the study were approved by the Institutional Animal Ethics Committee.

## Acknowledgments

We are grateful to Mr. Khem Singh Negi for genotyping mice and other technical help and to Mr. B. N. Roy for help with histopathology experiments. We thank Dr. Nagarajan for his help with animal experiments.

