## Supplementary figures for "VDR-calcium axis regulates the diet-driven metabolic shift during weaning"

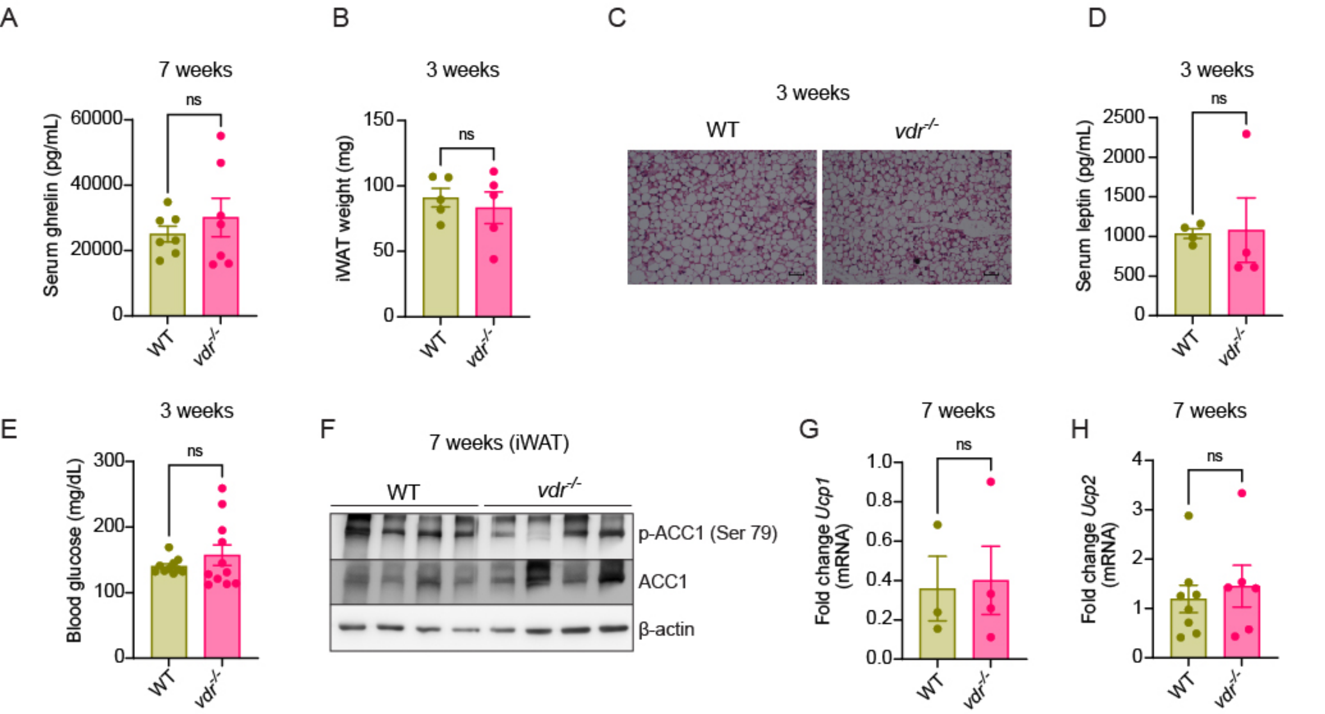


**Fig. S1. *vdr$^{-/-}$* mice exhibit severe energy deprivation and adipose atrophy post-weaning.** (A) Serum levels of ghrelin (in pg/mL) in 7-week WT and *vdr$^{-/-}$* mice under *ad libitum* fed state were measured using ELISA. (B) Inguinal WAT weight (in mg) in 3-week WT and *vdr$^{-/-}$* mice. (C) Representative images of H&E stained iWAT sections of 3-week WT and *vdr$^{-/-}$* mice. (D) Serum levels of leptin in 3-week WT and *vdr$^{-/-}$* mice, measured using ELISA under *ad libitum* fed state. (E) Blood glucose levels (in mg/dL) in 3-week WT and *vdr$^{-/-}$* mice under *ad libitum* fed state. (F) Western blot representing the levels of phospho-ACC1 (Ser 79) and total ACC1 in iWAT of 7-week WT and *vdr$^{-/-}$* mice. (G&H) mRNA levels of *Ucp1* (G) and *Ucp2* (H) in iWAT of 7-week WT and *vdr$^{-/-}$* mice as quantified using qRT-PCR. All graphs show mean ± SEM. Statistical significance was determined by unpaired t-test (*$p < 0.05$, **$p < 0.01$, ***$p < 0.001$ and ****$p < 0.0001$). The number of samples is denoted by the dots in the graphs.

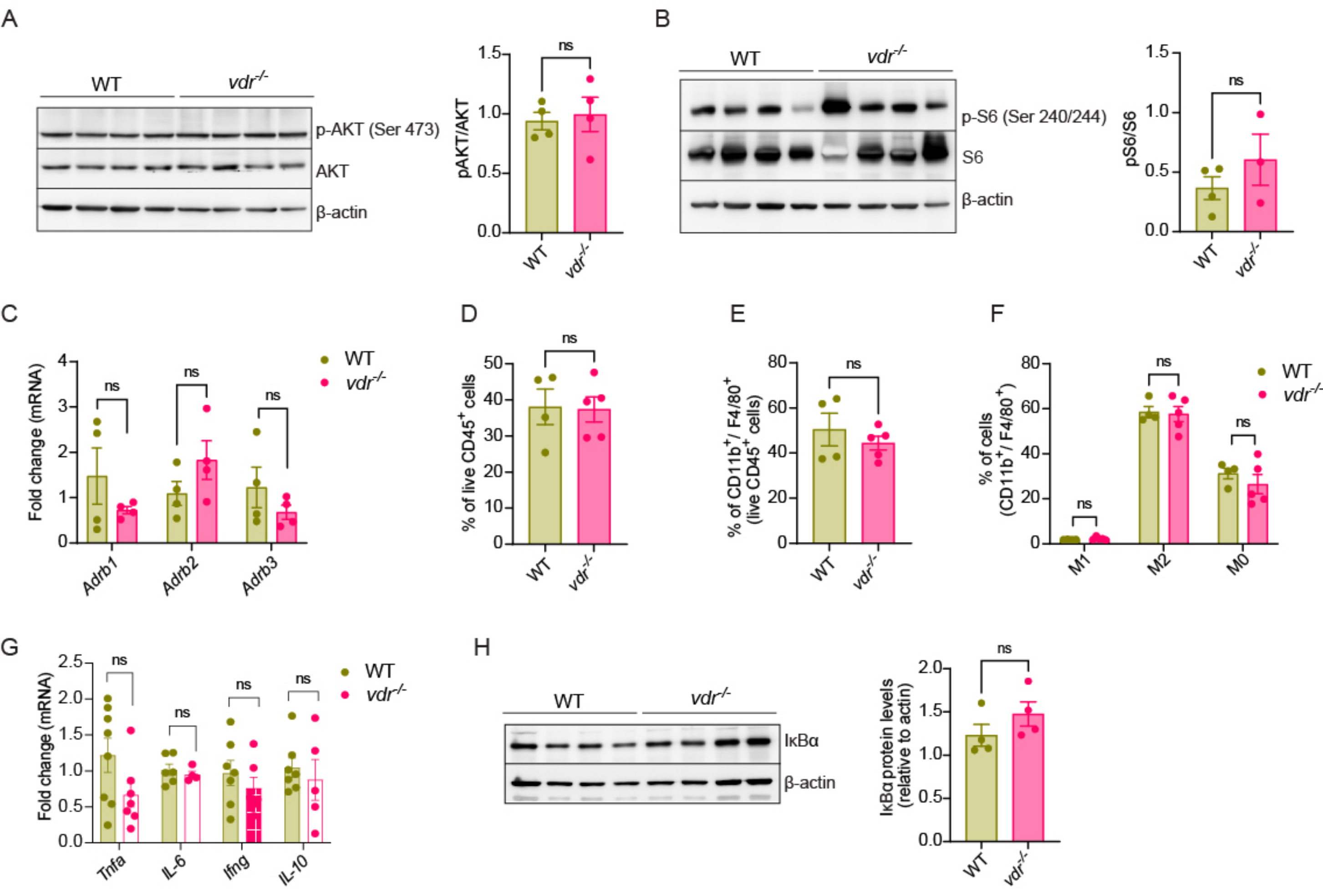


**Fig. S2. $vdr^{-/-}$ mice display increased norepinephrine levels with increased PKA and ERK activity in iWAT.** (A) Western blot representing the levels of phospho-Akt (Ser 473), total Akt, and the ratio of phospho-Akt to total Akt (right panel) in iWAT of 7-week WT and $vdr^{-/-}$ mice. (B) Western blot representing the levels of phospho-S6 (Ser 240/244), total S6, and the ratio of phospho-S6 to total S6 (right panel) in iWAT of 7-week WT and $vdr^{-/-}$ mice. (C) mRNA expression levels of β-adrenergic receptors 1, 2 and 3 in iWAT of 7-week WT and $vdr^{-/-}$ mice as quantified using qRT-PCR. (D-F) Quantification of the frequency of live CD45$^+$ cells (D), CD11b$^+$/F4/80$^+$ cells (E) and M1 (CD11c$^+$/CD206$^-$), M2 (CD11c$^-$/CD206$^+$) and M0 (CD11c$^-$/CD206$^-$) cells (F). (G) mRNA expression levels of *Tnfa*, *IL-6*, *Ifng* and *IL-10* in iWAT of 7-week WT and $vdr^{-/-}$ mice as quantified using qRT-PCR. (H) Western blot showing protein levels of IκBα and densitometric analysis (right panel) in iWAT of 7-week WT and $vdr^{-/-}$ mice. All graphs show mean ± SEM. Statistical significance was determined by unpaired t-test (*p < 0.05, **p < 0.01, ***p< 0.001 and ****p < 0.0001). The number of samples is denoted by the dots in the graphs.

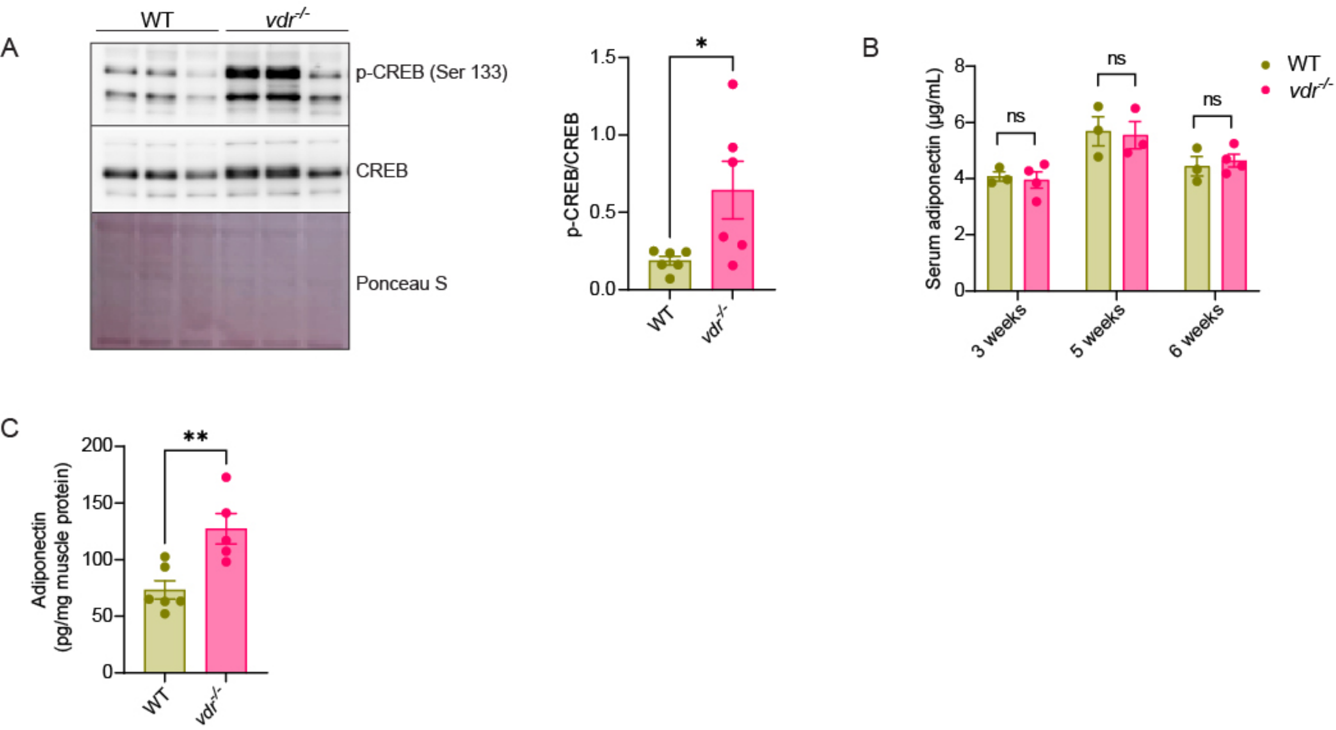


**Fig. S3. *vdr*$^{-/-}$ mice show increased basal mitochondrial activity in the adipose.** (A) Western blot showing the protein levels of p-CREB (Ser 133), total CREB and the ratio of phospho-CREB to total CREB in iWAT of 7-week WT and *vdr*$^{-/-}$ mice. Ponceau stained blot is given to show equal loading. (B) Serum adiponectin levels (in µg/mL) in WT and *vdr*$^{-/-}$ mice at 3, 5, and 6 weeks of age as measured using ELISA. (C) Muscle adiponectin levels (in pg/mg muscle protein) in 7-week WT and *vdr*$^{-/-}$ mice, measured using ELISA. All graphs show mean ± SEM. Statistical significance was determined by unpaired t-test (*p < 0.05, **p < 0.01, ***p< 0.001 and ****p < 0.0001). The number of samples is denoted by the dots in the graphs.

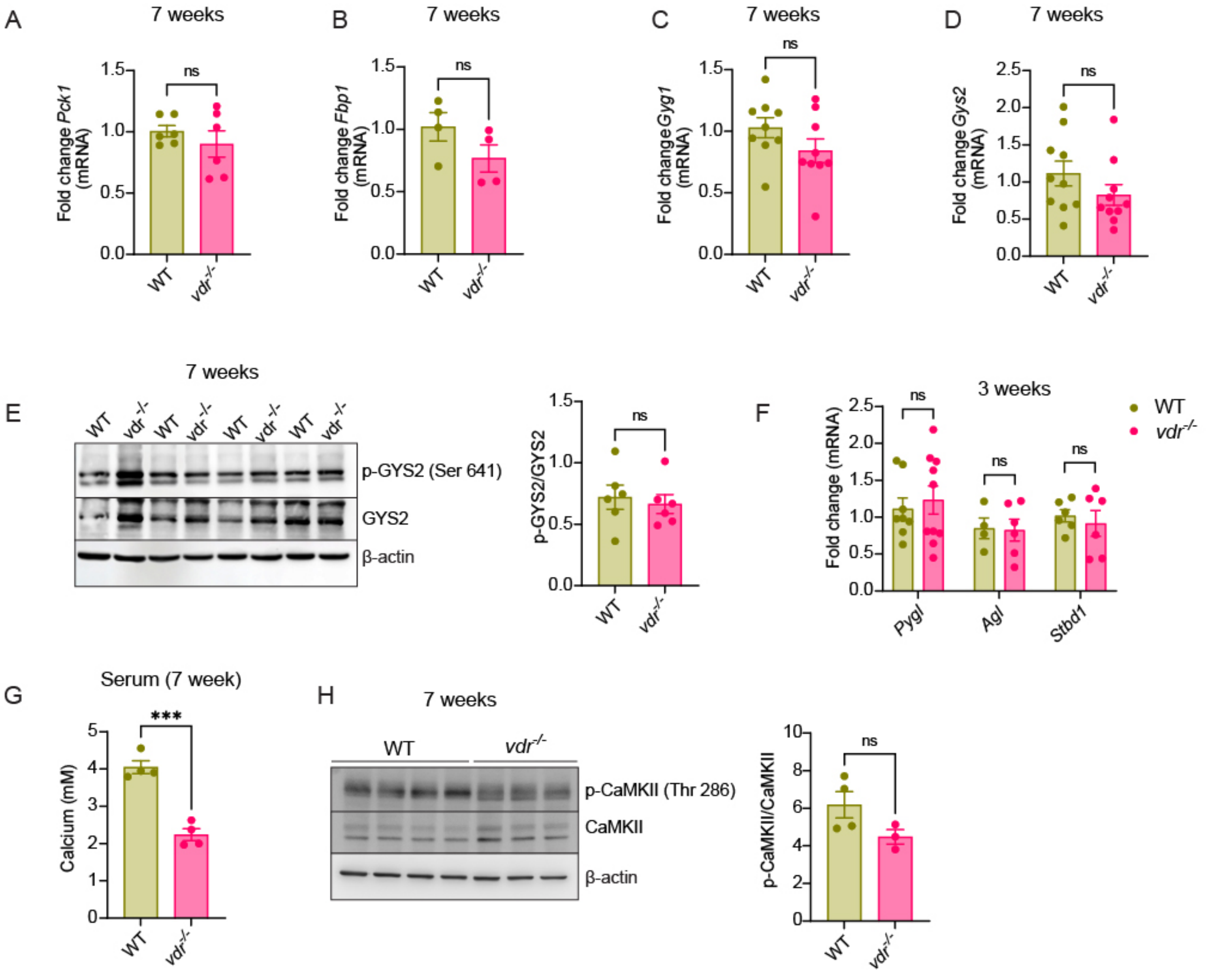


**Fig. S4. $vdr^{-/-}$ mice show defective liver glycogenolysis and increased protein phosphatase 1 level.** (A-D) mRNA levels of *Pck1* (A), *Fbp1* (B), *Gyg1* (C), and *Gys2* (D) in the liver of 7-week WT and $vdr^{-/-}$ mice as quantified using qRT-PCR. (E) Western blot representing the levels of phospho-GYS2 (Ser 641), total GYS2, and the ratio of phospho-GYS2 to total GYS2 (right panel) in liver (*ad libitum* fed state) of 7-week WT and $vdr^{-/-}$ mice. (F) mRNA levels of glycogen phosphorylase (*Pygl*), glycogen debranching enzyme (*Agl*) and starch binding domain-containing protein 1 (*Stbd1*) in the liver of 3-week WT and $vdr^{-/-}$ mice as quantified using qRT-PCR. (G) Serum calcium levels (in mM) of 7-week WT and $vdr^{-/-}$ mice, measured using ELISA. (H) Western blot showing protein levels of phospho-CaMKII (Thr 286), total CaMKII and the ratio of phospho-CaMKII to total CaMKII (right panel) in liver (12-hour fasted state) of 7-week WT and $vdr^{-/-}$ mice. All graphs show mean ± SEM. Statistical significance was determined by unpaired t-test (*$p < 0.05$, **$p < 0.01$, ***$p < 0.001$ and ****$p < 0.0001$). The number of samples is denoted by the dots in the graphs.

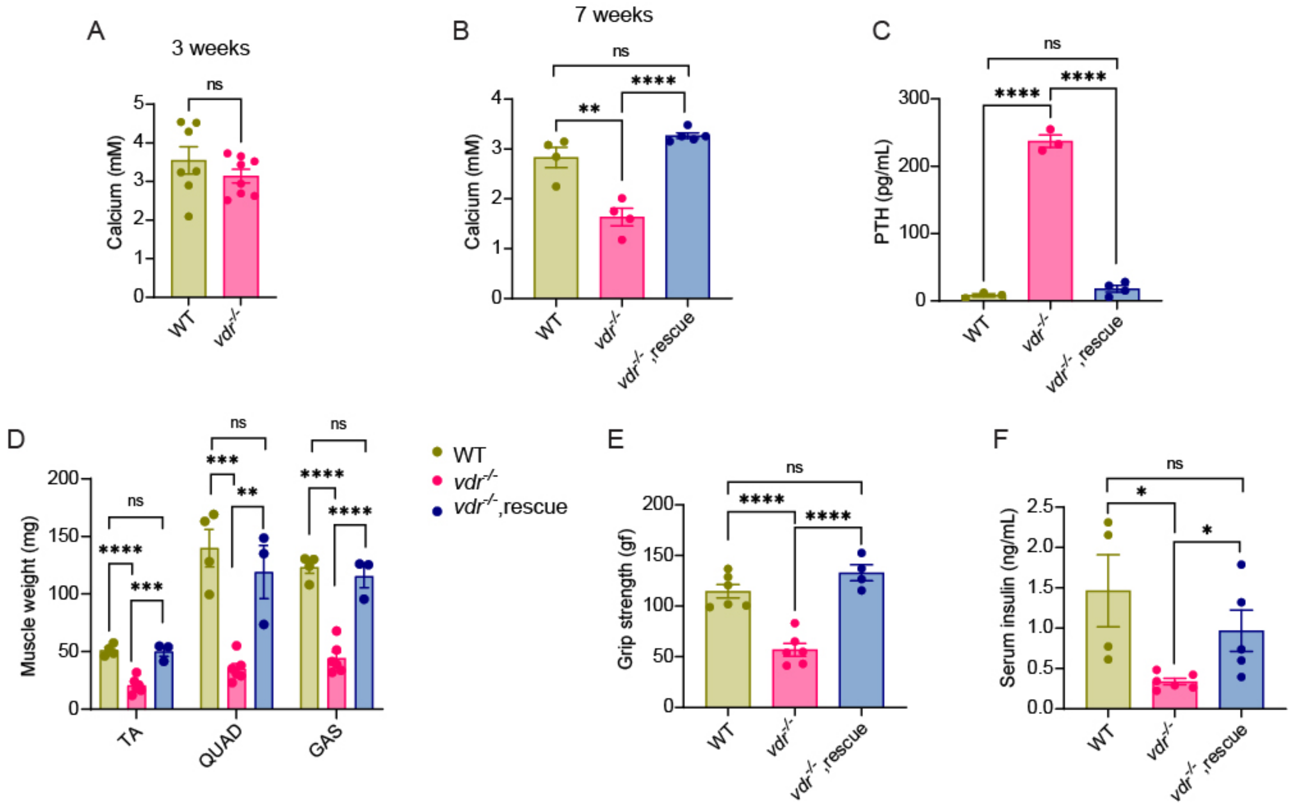


**Fig. S5. Weaning on a calcium-rich diet prevents adipose wasting and rescues glycogenolysis defect in *vdr*$^{-/-}$.** (A) Serum calcium levels (in mM) of 3-week WT and *vdr*$^{-/-}$ mice, measured using ELISA. (B) Serum calcium levels (in mM) of 7-week WT and *vdr*$^{-/-}$ mice on chow and rescue diet, measured using ELISA. (C) Serum PTH levels (in pg/mL) of 7-week WT and *vdr*$^{-/-}$ mice on chow and rescue diet measured using ELISA. (D) TA, QUAD, GAS muscle weight of 7-week-old WT and *vdr*$^{-/-}$ mice on chow and rescue diet. (E) Forelimb grip strength (in gF) of 7-week WT and *vdr*$^{-/-}$ mice on chow and rescue diet. (F) Serum insulin levels (in ng/mL) of 7-week WT and *vdr*$^{-/-}$ mice on chow and rescue diet measured using ELISA. All graphs show mean ± SEM. Statistical significance was determined by unpaired t-test (*$p < 0.05$, **$p < 0.01$, ***$p < 0.001$ and ****$p < 0.0001$). The number of samples is denoted by the dots in the graphs.
